# Genome sequencing sheds light on the contribution of structural variants to *Brassica oleracea* diversification

**DOI:** 10.1101/2020.10.15.340224

**Authors:** Ning Guo, Shenyun Wang, Lei Gao, Yongming Liu, Mengmeng Duan, Guixiang Wang, Jingjing Li, Meng Yang, Mei Zong, Shuo Han, Yanzheng Pei, Theo Borm, Honghe Sun, Liming Miao, Di Liu, Fangwei Yu, Wei Zhang, Heliang Ji, Chaohui Zhu, Yong Xu, Guusje Bonnema, Jianbin Li, Zhangjun Fei, Fan Liu

**Affiliations:** National Engineering Research Center for Vegetables, Beijing Academy of Agriculture and Forestry Sciences, Key Laboratory of Biology and Genetic Improvement of Horticultural Crops (North China), Beijing Key Laboratory of Vegetable Germplasm Improvement, Beijing, China; Jiangsu Key Laboratory for Horticultural Crop Genetic Improvement, Vegetable Research Institute, Jiangsu Academy of Agricultural Science, Nanjing, Jiangsu, China; Boyce Thompson Institute, Ithaca, NY, USA; Plant Breeding, Wageningen University and Research, Wageningen, The Netherlands; Nextomics Biosciences Institute, Wuhan, China; CAS Key Laboratory of Plant Germplasm Enhancement and Specialty Agriculture, Wuhan Botanical Garden, Chinese Academy of Sciences, Wuhan, Hubei, China; Tianjin GengYun Seed Company, Tianjin, China; Fuzhou Institute of Vegetable Science, Fuzhou, Fujian, China; US Department of Agriculture-Agricultural Research Service, Robert W. Holley Center for Agriculture and Health, Ithaca, NY, USA

## Abstract

*Brassica oleracea* includes several morphologically diverse, economically important vegetable crops. Here we present high-quality chromosome-scale genome assemblies for two *B. oleracea* morphotypes, cauliflower and cabbage. Direct comparison of these two assemblies identifies ~120 K high-confidence structural variants (SVs). Population analysis of 271 *B. oleracea* accessions using these SVs clearly separates different morphotypes, suggesting the association of SVs with *B. oleracea* intraspecific divergence. Genes affected by SVs selected between cauliflower and cabbage are enriched with functions related to response to stress and stimulus and meristem and flower development. Furthermore, genes affected by selected SVs and involved in the switch from vegetative to generative growth that defines curd initiation, inflorescence meristem proliferation for curd formation, maintenance and enlargement, are identified, providing insights into the regulatory network of curd development. This study reveals the important roles of SVs in diversification of different morphotypes of *B. oleracea*, and the newly assembled genomes and the SVs provide rich resources for future research and breeding.

## Introduction

*Brassica oleracea* includes several diverse dominant vegetable crops with a worldwide total production of nearly 100 million tons in 2018 (http://www.fao.org/faostat). The extreme diversity of this species is unique with morphotypes selected for the enlargement of distinct organs that represent the harvested product, e.g., inflorescences for cauliflower *(B. oleracea* var. *botrytis)* and broccoli (*B. oleracea* var. *italica)*, leafy heads (terminal leaf bud) for cabbage (*B. oleracea* var. *capitata)*, lateral leaf buds for Brussels sprouts (*B. oleracea* var. *gemmifera)*, leaves for kale (*B. oleracea* var. *alboglabra)* and tuberous stems for kohlrabi (*B. oleracea* var. *gongylodes)* (Dixon, 2007; Cheng et al., 2014). Reference genome sequences have been generated for different morphotypes of *B. oleracea* during the past several years, including kale (Parkin et al., 2014), cabbage (Liu et al., 2014; Cai et al., 2020; Lv et al., 2020), cauliflower (Sun et al., 2019) and broccoli (Belser et al., 2018). These genome sequences have greatly facilitated genetic variant analyses for a better understanding of the genetic diversity, population structure, and evolution and domestication of *B. oleracea.*

Structural variants (SVs) including insertions, deletions, duplications and translocations are abundant throughout plant genomes and are more likely to cause phenotype changes than single nucleotide polymorphisms (SNPs) (Zhou et al., 2019; Song et al., 2020). Numerous SVs have been identified as causal genetic variants for important agronomic traits of various crops, such as the 4.7-kb insertion into the third exon of the *Or* gene leading to the orange curd in cauliflower (Lu et al., 2006), the 3.7-kb insertion in the upstream region of *BnaA9.CYP78A9* leading to the long siliques and large seeds of *Brassica napus* (Shi et al., 2019), and the 621-bp insertion in the promoter region of *BnaFLC.A10* contributing to the adaptation of rapeseed to winter cultivation environments (Yin et al., 2020). Previous genome-wide variant analyses in *B. oleracea* focused on SNPs and small indels (Cheng et al., 2016; Stansell et al., 2018) with genomic SVs largely ignored, mainly due to the limitations of using short sequencing reads in genetic variant identification. SV calling through mapping short sequencing reads to a reference genome is subject to high levels of both false negatives and false positives (Sedlazeck et al., 2018), especially for highly repetitive plant genomes such as those of *B. oleracea*. Therefore, to date population dynamics of SVs in different *B. oleracea* morphotypes remain largely unexplored.

Recently, approaches by direct comparison of high-quality chromosome-level genome assemblies and/or mapping long reads generated using PacBio or Nanopore sequencing technologies to reference genomes have proven to be highly accurate for SV detection in large and complex plant genomes (Alonge et al., 2020; Liu et al., 2020). In this study, we generated high-quality chromosome-scale genome assemblies for both cauliflower and cabbage using PacBio long reads and the high-throughput chromosome conformation capture (Hi-C) technology. Through direct genome comparison combined with long read mapping, we identified a total of 119,156 high-confidence SVs between these two genomes. We further generated and collected genome resequencing data of 271 *B. oleracea* accessions belonging to different morphotypes, and these data were used to genotype the 119,156 high-confidence SVs in these accessions. Allele frequencies of these SVs were investigated in different *B. oleracea* morphotypes, and mainly compared between cauliflower and cabbage populations. Together with gene expression analysis, we demonstrated the contribution of SVs to the regulation of cauliflower curd formation.

## Results

### *De novo* assembly of cauliflower and cabbage genomes

The inbred lines cauliflower Korso_1401 (hereafter Korso) and pointed cabbage OX-heart_923 (hereafter OX-heart) were selected for genome sequencing (**Supplementary Fig. 1**). Approximately 70.0 Gb PacBio sequences were generated for each accession, covering about 120× of the Korso and OX-heart genomes, which had estimated sizes of 566.9 Mb and 587.7 Mb, respectively (**Supplementary Fig. 2**). These PacBio reads were *de novo* assembled into contigs and errors in the assembled contigs were corrected using both PacBio long reads and Illumina short reads (~100 Gb for each accession). In addition, a genome map was assembled from 242.2 Gb cleaned BioNano optical map data for Korso and used to connect the assembled contigs. Furthermore, 285.8 and 453.0 million cleaned Hi-C read pairs, among which 58.6 and 140.0 million were valid, were used for pseudochromosome construction for Korso and OX-heart, respectively. The final genome assemblies of Korso and OX-heart comprised 615 and 973 contigs, respectively, with cumulative lengths of 549.7 Mb and 565.4 Mb, and N50 sizes of 4.97 Mb and 3.10 Mb (**Supplementary Table 1**). A total of 544.4 Mb and 539.1 Mb, accounting for 99.0% and 95.3% of the Korso and OX-heart assemblies, respectively, were clustered into nine pseudomolecules. The Hi-C heatmaps (**Supplementary Fig. 3**) and the good synteny between Korso and OX-heart assemblies and the broccoli HDEM assembly (Belser et al., 2018) (**Supplementary Fig. 4**) supported their chromosome-scale structures.

Around 99.8% of the Illumina genomic reads could be mapped back to the Korso and OXheart assemblies, with 99.6% of the assemblies covered by at least 5 reads. Based on the alignments, the estimated base error rates of the Korso and OX-heart assemblies were 1.23×10^-5^ and 5.6×10^-5^, respectively (**Supplementary Table 2**). BUSCO analysis (Waterhouse et al., 2018) showed that 97.2% and 96.5% core conserved plant genes were completely assembled in Korso and OX-heart. In addition, up to 98.0% of the RNA-Seq reads could be mapped to the assemblies (**Supplementary Table 3**). Together, these results demonstrated the high quality of the Korso and OX-heart assemblies.

### Genome annotation and comparative genomics

Approximately 60.7% and 62.0% of the Korso and OX-heart assemblies were annotated as repetitive elements, respectively, with the *Gypsy-* and *Copia*-like retrotransposons representing the most abundant families in both genomes (**Supplementary Table 4**). Full-length long terminal repeat retrotransposons (LTR-RTs) were then extracted from the Korso, OX-heart and *B. rapa* (V3.0) (Zhang et al., 2018) genomes (**Supplementary Table 5**). Insertion time estimation of these intact LTR-RTs unraveled two LTR-RT bursts that occurred in Korso and OX-heart, around 0.2 and 1.5 million years ago (mya), respectively (**Supplementary Fig. 5**). In contrast, in *B. rapa*, most of the LRT-RT formed recently, with more than 30% of the identified intact LTR-RTs younger than 0.2 mya, compared to 16.3% and 15.9% in Korso and OX-heart, respectively.

The high-quality Korso and OX-heart assemblies allowed us to precisely identify the centromere locations. The determined positions of centromeres on each chromosome in both genomes (**Supplementary Fig. 6**) were consistent with the previously determined centromere locations using fluorescent in situ hybridization (FISH) analysis (Xiong and Pires, 2011). As expected, repetitive elements were enriched in the centromere regions. Different repeat families displayed clearly different patterns on the chromosomes, e.g., *Copia*-type LTRs were mainly in centromeres, while *Gypsy*-type LTRs were in pericentromeric regions (Fig. 1a).

**Fig. 1.**
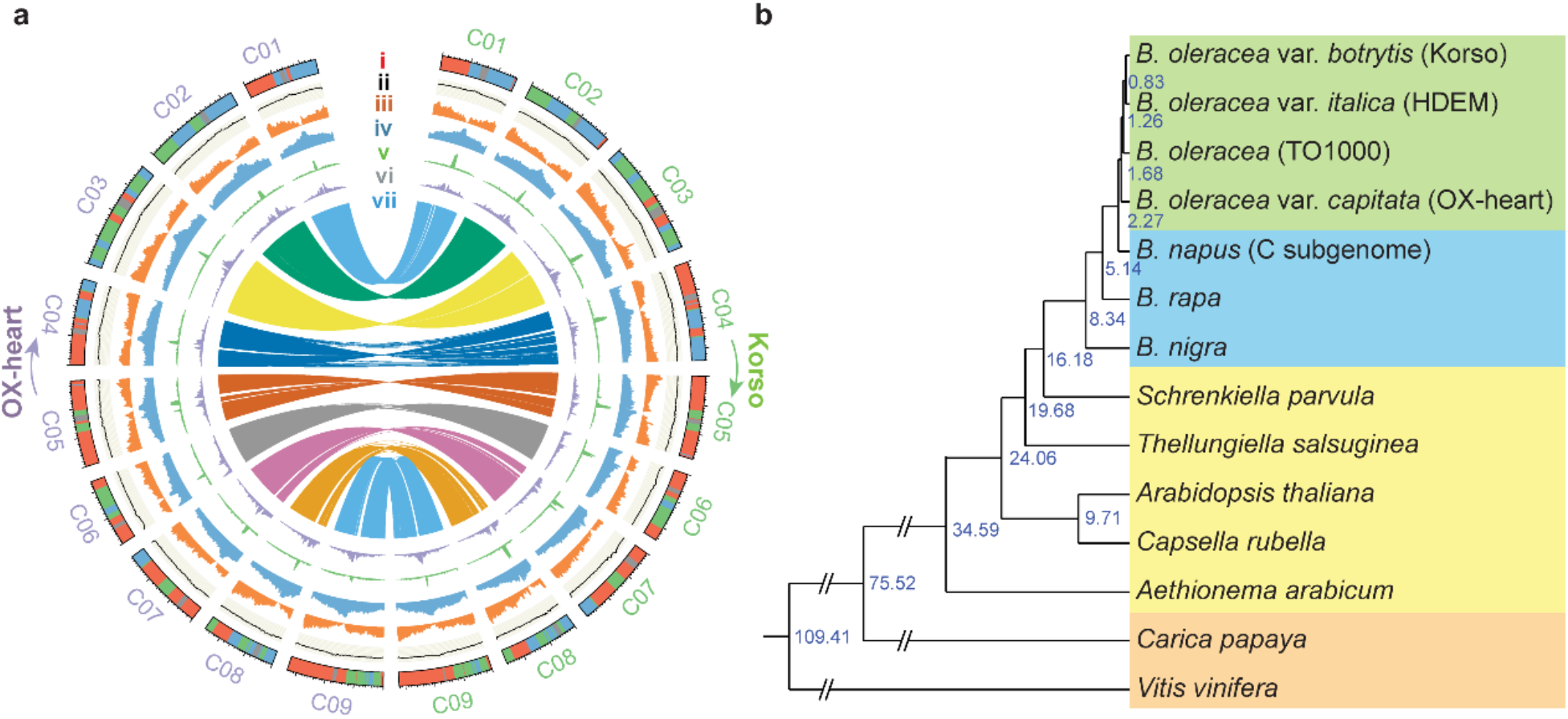
Genomes of cauliflower Korso and point cabbage OX-heart. **a**, Features of the Korso and OX-heart genomes. i, Ideogram of the chromosomes. Red, green, blue and grey colors indicate the LF, MF1, MF2 subgenomes and centromere regions, respectively. ii. GC content. iii, Gene density. iv, Repeat density. v, *Copia*-type LTR density. vi, Gypsy-type LTR density. vii, Synteny blocks between Korso and OX-heart genomes. **b**, Phylogenetic tree of 14 plant species/varieties and their estimated divergence times (million years ago) based on 1,638 single-copy orthologous genes.

A total of 60,640 and 62,232 high-confidence protein-coding genes were predicted from Korso and OX-heart genomes, respectively, using an integrated strategy combining *ab initio*, transcript-based and homology-based predictions. Among these predicted genes, 70.9% and 76.4% were supported by transcriptome evidence, and 91.0% and 90.0% had homologs in other plant species. BUSCO analysis (Waterhouse et al., 2017) revealed the completeness of 96.1% and 94.9% of Korso and OX-heart predicted genes, respectively.

Synteny analysis of Korso, OX-heart, *B. rapa* and *A. thaliana* genomes confirmed the whole genome triplication (WGT) and subsequent sub-genome divergence in *Brassica* species (Wang et al., 2011; Liu et al., 2014; Cheng et al., 2016) (**Supplementary Figs. 7** and **8**). Based on these syntenic relationships, we identified the triplicated regions within Korso and OX-heart genomes and divided them into three subgenomes based on their retained gene densities (Fig. 1). As previously reported in *B. rapa* (Wang et al., 2011) and *B. oleracea* (Liu et al., 2014), the three subgenomes of Korso and OX-heart, LF (the least fractionated), MF1 (the medium fractionated) and MF2 (the most fractionated), showed the same biased retention pattern of duplicated genes during diploidization (Xie et al., 2019) (**Supplementary Fig. 8a,b**). Duplicated gene copies left in different subgenomes displayed diverged gene expression patterns, with the copies located in LF generally having higher expression levels than those in MF1 and MF2 (**Supplementary Fig. 8c,d**).

We compared protein sequences of predicted genes from four *B. oleracea* accessions (cauliflower Korso, pointed cabbage OX-heart, broccoli HDEM and kale like rapid cycling TO1000), three other *Brassica* species, *B. rapa, B. nigra*, and the C subgenome of *B. napus*, five other Brassiaceae species (*Aethionema arabicum*, *Arabidopsis thaliana*, *Capsella rubella*, *Thellungiella salsuginea* and *Schrenkiella parvula*), and two outgroups (grape and papaya). A phylogenetic tree was constructed using 1,638 single-copy orthologous genes, which indicated that cabbage and the common ancestor of cauliflower and broccoli diverged about 1.68 mya, the extant *B. oleracea* and the donor of *B. napus* C subgenome diverged about 2.27 mya, and *Brassica* diverged from other Brassiaceae species about 16.18 mya (Fig. 1b).

### SVs between genomes of Korso and OX-heart

By taking advantage of the high-quality genome assemblies of Korso and OX-heart, we were able to identify high-confidence SVs through direct genome comparison combined with PacBio long read mapping. The Korso and OX-heart assemblies displayed very high collinearity indicating the balanced rearrangements (inversions and translocations) were not profound between them (**Supplementary Fig. 4**). Therefore, in this study, we focused on the unbalanced SVs, mainly indels. A total of 119,156 SVs were identified between genomes of Korso and OX-heart, with sizes ranging from 10 bp to 667 kb with a clear bias to the relatively short ones (**Supplementary Table 6** and **Supplementary Fig. 9**).

SVs in gene bodies and promoter regions can affect the function or expression of the corresponding genes. The SV regions accounted for 14.5% and 15.0% of the total genome sizes of Korso and OX-heart, 10.0% and 11.3% of the gene regions, and 5.9% and 6.6% of the coding sequences, respectively, suggesting a functional constraint against the occurrence of SVs in genes, especially in coding regions, while no obvious restriction of SVs in promoter regions was detected (**Supplementary Table 7**). More than half of the annotated genes in Korso (58.5%) and OX-heart (58.6%) were affected by at least one SV in their gene bodies or promoter regions, with a functional enrichment in diverse biological processes, such as cellular component organization, response to stress and stimulus, signal transduction, cell differentiation, embryo development, gene expression and epigenetic regulation, and flower and meristem development (**Supplementary Fig. 10**). We detected several previously described SVs in *B. oleracea*, including the two indels in *BoFLC3* related to subtropical adaptation of broccoli (Lin et al., 2018), and the two indels in *BoFRIa* related to winter annual or biennial habit of cauliflower and cabbage (Irwin et al., 2012).

### Population dynamics of SVs in different *B. oleracea* morphotypes

Cabbage and cauliflower represent two extreme morphotypes of the *B. oleracea* species, and identifying genomic variations underlying the formation of their unique phenotypes (e.g., leafy head and curd) would provide novel insights into the molecular regulation of these important traits as well as important information for facilitating breeding. The high-quality SVs that we identified between Korso and OX-heart provided a valuable reference to investigate their dynamics in different morphologically diverged *B. oleracea* accessions. For this purpose, we performed genome resequencing of 163 *B. oleracea* accessions, including 89 cauliflower, 65 cabbage and 9 broccoli accessions. We also collected resequencing data of an additional 108 *B. oleracea* accessions reported in Cheng et al. (2016), including 15 cauliflower, 39 cabbage, 24 broccoli, 18 kohlrabi, four Chinese kale, two curly kale, two kale, two Brussels sprout and two wild *B. oleracea* accessions (**Supplementary Table 8**). Among these 271 accessions, 211 were sequenced to a depth of more than 10×. The 119,156 high-quality reference SVs were genotyped in these 271 accessions based on the alignments of genome sequencing reads to the Korso and OX-heart genomes. To assess the accuracy of our SV genotyping, we genotyped the reference SVs in Korso and OX-heart by mapping their Illumina short reads to both these genomes, respectively. More than 86% of SVs could be genotyped, while only 0.1% were falsely genotyped, suggesting high sensitivity and accuracy of our genotyping. The SV genotyping rate in each accession ranged from 41.3% to 80.2%, with 187 (69.0%) and 254 (93.7%) accessions having a genotyping rate greater than 70% and 60%, respectively (**Supplementary Fig. 11** and **Supplementary Table 8**). In total, 89,882 (75.4%) SVs were successfully genotyped in more than 50% of the 271 accessions.

SV allele frequency variations among different groups of *B. oleracea* are mainly a result of domestication for different desirable traits and adaptation to different environments. As expected, SV loci with the homozygous Korso alleles were prevalent in cauliflower accessions, taking up an average of 82.3% of the genotyped SVs in each accession, whereas in cabbage accessions, the homozygous OX-heart alleles were prevalent, with an average frequency of 61.7% (Fig. 2a and **Supplementary Table 8**). Phylogenetic and principal component analyses (PCA) using the SVs clearly divided cauliflower, cabbage, broccoli and kohlrabi accessions into different groups (Fig. 2b and 2c), which were concordant with the patterns revealed by SNP data in our analysis based on the same 271 accessions (**Supplementary Fig. 12**) and the previous report based on 119 accessions (Cheng et al., 2016), further supporting that our SV detection and genotyping were highly reliable.

**Fig. 2.**
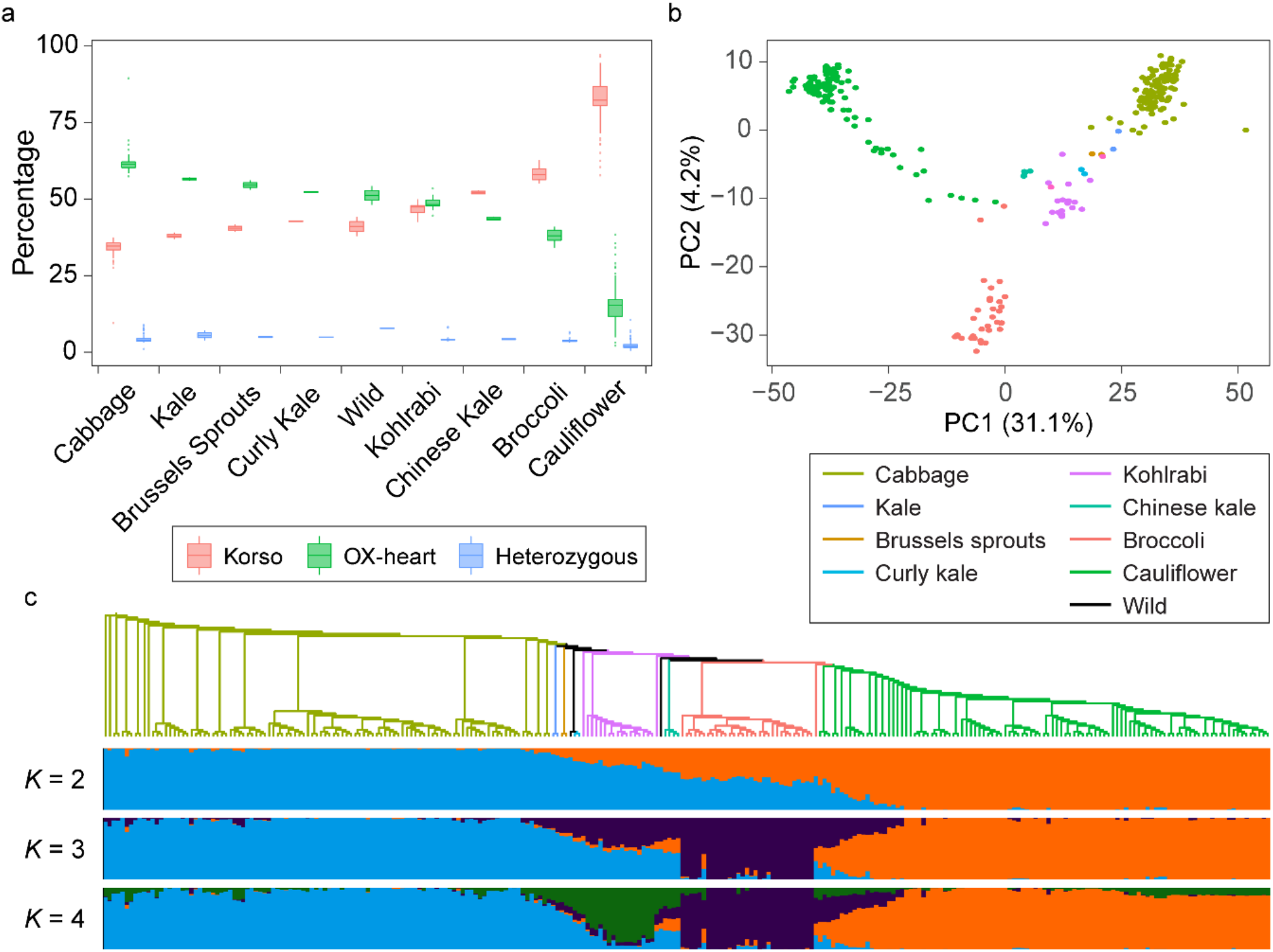
SVs in different *B. oleracea* morphotypes. **a**, Percentages of SVs with different genotypes in accessions of different morphotypes. **b**, Principal component analysis of *B. oleracea* accessions based on SVs. **c**, Maximum-likelihood tree and model-based clustering of the 271 *B. oleracea* accessions using SVs. Branch colors of the tree indicate different morphotypes as in **b**. *K*, number of ancestral kinships.

To identify SVs potentially related to the specific traits of cauliflower or cabbage, we extracted a total of 49,904 SVs with significantly different allele frequencies between cauliflower and cabbage populations (Fig. 3a). Among these SVs, 49,285 (98.8%) had significantly higher allele frequencies of Korso genotypes in cauliflowers than in cabbages, while only 550 represented higher OX-heart allele frequencies in cauliflowers than in cabbages. These potentially selected SVs were distributed across the chromosomes without conspicuous hotspots (**Supplementary Fig. 13**). Such prevalence of selected SVs across the genome is consistent with the relatively large divergence time (~1.68 mya) between the two highly specialized *B. oleracea* morphotypes and their independent evolution and domestication history (Fig. 1b).

**Fig. 3.**
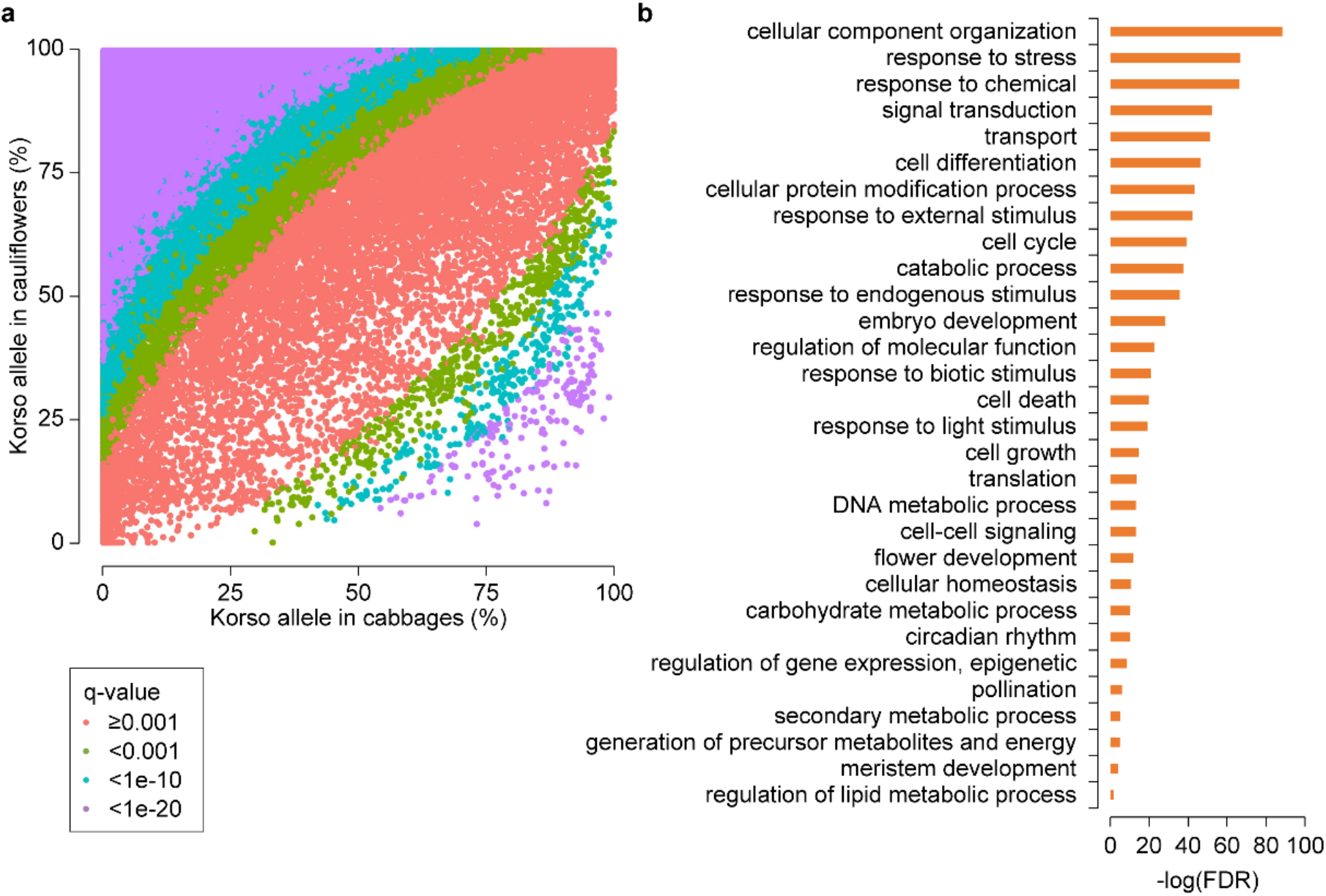
SV divergence between cauliflower and cabbage. **a**, Scatter plot showing Korso-allele frequencies of SVs in cabbage and cauliflower groups. q-value, Bonferroni-corrected p value of Fisher’s exact test. **b**, GO terms enriched in genes overlapping with SVs with significant allele frequency differences between cabbage and cauliflower groups.

In Korso and OX-heart genomes, 21,111 and 21,400 genes, respectively, overlapped with at least one selected SV in their gene bodies or promoter regions, with 6,059 and 6,344 overlapping with selected SVs in CDS regions. GO enrichment analyses of these genes with selected SVs revealed that those related to signal transduction, response to stimulus, cell differentiation, cell cycle, embryo development, cell growth and cell death, and flower development were significantly overrepresented (Fig. 3b), showing potential associations with the distinct phenotypes of cauliflower (curd) and cabbage (leafy head).

### Selected SVs provide insights into the evolution of cauliflower curd formation

The curd of cauliflower is composed of a spirally iterative pattern of primary inflorescence meristems with floral primordia arrested in their development (Sadik, 1962; Kieffer et al., 1998). The first insight in genetic control of the curd-like structure was achieved through characterization of the *Arabidopsis ap1* and *cal* double mutant with a cauliflower curd phenotype (Kempin et al., 1995). Subsequently, several studies indicated that the genetic nature of the cauliflower curd appears more complex (Smith and King, 2000; Labate et al., 2006; Duclos and Björkman, 2008). Here, we retrieved a total of 294 genes harboring selected SVs in their promoters or gene regions and whose homologues in *Arabidopsis* have been reported to function in flowering time and floral development, meristem maintenance and determination, organ size control, and shoot or inflorescence architecture (**Supplementary Table 9**). In addition, RNA-Seq analysis of five stages from vegetative shoot apical meristem (SAM) to enlarged curd was conducted to reveal the potential roles of SVs in curd formation and development.

#### Transition from vegetative to generative development

The first stage of curd initiation corresponds with the switch from vegetative to generative development (Fig. 4a). Timely transition to the generative stage in cauliflower is essential for curd formation, while for cabbage a prolonged vegetative stage is needed for the proper development of the leafy head. The MADS box transcription factor FLC, a flowering time integrator in the vernalization and autonomous pathways, acts as a repressor of flowering (Michaels and Amasino, 1999; Sheldon et al., 1999). Several studies have demonstrated the roles of *FLC* paralogues in flowering time in diverse *B. oleracea* morphotypes (Okazaki et al., 2007; Razi et al., 2008; Ridge et al., 2015; Irwin et al., 2016; Lin et al., 2018). A 3,371-bp insertion (SV_b_92666a) in the promoter of *BoFLC1.1* in Korso was found under strong differential selection, present in 99% and 88% of the cauliflower and broccoli accessions, respectively, while only in 9% of cabbage accessions (Fig. 4b). *BoFLC1.1* and its two tandem paralogs (*BoFLC1.2* and *BoFLC1.3*), as well as *BoFLC3* were all significantly down-regulated at the transition stage (Fig. 4c). The Korso allele of *BoFLC3* contains a 263-bp deletion (SV_w_24534) and a 49-bp insertion (SV_w_24533) in the first intron. The effect of the structure of the *FLC* first intron on flowering time has been reported in *Arabidopsis* and cruciferous crops (Sheldon et al., 2002; Lin et al., 2018; Wang et al., 2018). We found that at these two SV loci, the Korso alleles were predominant in cauliflower (86.7% and 86.4%) and broccoli (96.9% and 92.9%), but rare in the cabbage accessions (9.7% and 8.7%) (Fig. 4b).

**Fig. 4.**
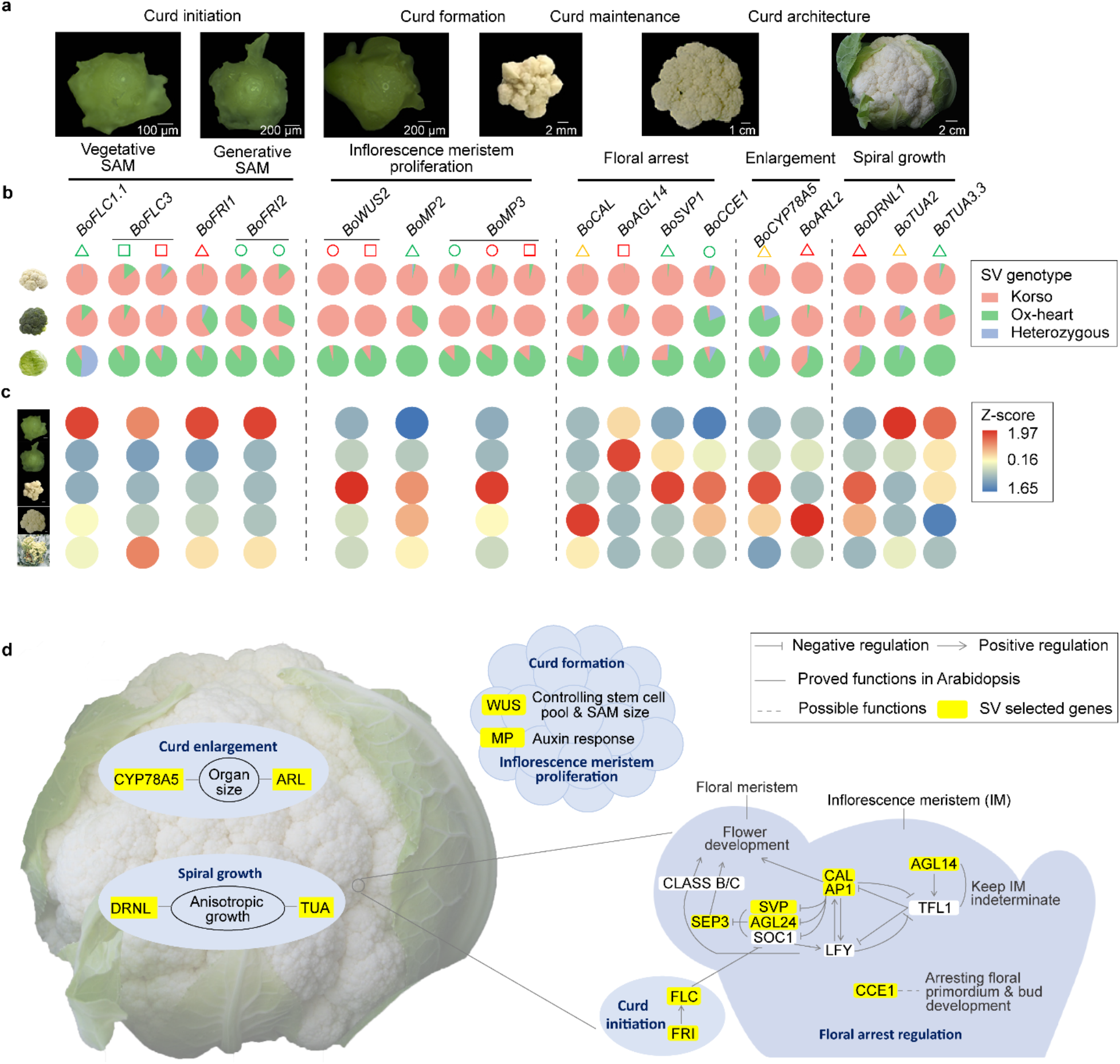
SV contribution to cauliflower curd formation. **a**, SAM and curd at different stages of curd development. **b**, Korso-allele frequencies of SVs overlapping with candidate genes in cauliflower, broccoli and cabbage. Triangles, circles, and squares indicate SVs in promoter, CDS and intron regions, respectively, and their different colors, green, red and yellow, indicate different types of SVs, insertion, deletion and substitution, respectively, in Korso compare to OX-heart. **c**, Heatmaps showing expression of candidate genes at different stages of curd development. The expression values (FPKM) were normalized by Z-score. **d**, Proposed regulatory network for curd formation and development of cauliflower. Genes with yellow background had selected SVs between cauliflower and cabbage.

The FLC function is activated by FRI (Geraldo et al., 2009), which has been identified as a candidate gene in the QTL region for temperature-dependent timing of curd induction in cauliflower (Hasan et al., 2016). Two *FRI* homologues, *BoFRI1* and *BoFRI2*, were identified in both Korso and OX-heart genomes. A 743-bp deletion (SV_b_96002) in the promoter region of *BoFRI1* characterized the Korso allele. Most of the cauliflowers (98.0%) contained the homozygous Korso genotype, while the majority of cabbages (87.0%) harbored the homozygous OX-heart genotype (Fig. 4b). For *BoFRI2*, two insertions (12- and 21-bp, SV_w_31837 and SV_w_31838) were identified in its coding region, both displaying significant differences of genotype frequencies between cauliflower and cabbage (Fig. 4b). These two indels have been found to be related to winter annual or biennial habit of cauliflower and cabbage (Irwin et al., 2012). FES and SUF can form a putative transcription activator complex with FRI to promote *FLC* expression (Michaels et al., 2004; Choi et al., 2011). The *B. oleracea* homologues *BoFES1.1* and *BoSUF4.2* harbored selected SVs in cauliflowers compared to cabbages, and their expression was significantly down-regulated from the vegetative to the transition stage in cauliflower, similar to that of *BoFLC1s* and *BoFLC3* (**Supplementary Table 9**). Other genes involved in regulating *FLC* expression, including those involved in epigenetic modification such as the PRC1 and PRC2 complex components *BoVIN3, BoVIL2.3, BoVRN1.1*, also harbored selected SVs (**Supplementary Table 9**). Together these results suggested that the FLC-related autonomous and vernalization pathways might be affected by the differential SVs between cauliflower and cabbage, contributing to their different timing of switch to the generative stage.

#### Inflorescence meristem proliferation

The main process following curd initiation is the continuously regular spiral proliferation of undetermined inflorescence meristems that form the curd. Stem cell maintenance and meristem proliferation play key roles in this process. WUSCHEL acts as an auxin response rheostat to maintain apical stem cells in *Arabidopsis* (Ma et al., 2019). We identified a 12-bp in-frame deletion (SV_w_83072) in the second exon and a 21-bp insertion (SV_w_83073) in the first intron of *BoWUS2* in Korso. All sampled cauliflower and broccoli accessions had the homozygous Korso genotypes for both SVs, while the Korso alleles were rare (4%) in cabbage (Fig. 4b). The expression of *BoWUS2* was significantly up-regulated from vegetative to curd formation, with the highest expression at the curd formation stage (Fig. 4c), implying that these two SVs could play roles in the curd formation.

MP/ARF5 together with ANT and AIL play key roles in auxin-dependent organ initiation and phyllotactic patterning (Yamaguchi et al., 2013; Bhatia and Heisler, 2018). Selected SVs in promoters and gene regions of their homologues in *B. oleracea, BoMP2, BoMP3, BoANT, BoAIL5, BoAIL6* and *BoAIL7*, were identified (**Supplementary Table 9**). A 23-bp insertion (SV_w_71238) in the promoter of *BoMP2* in Korso is under strong selection in cauliflower (96.1% and 0% in cauliflower and cabbage accessions, respectively). An 11-bp insertion and a 23-bp deletion (SV_w_92482 and SV_w_92481) in the CDS and a 14-bp deletion (SV_w_92433) in the intron of *BoMP3* in Korso are under strong selection in cauliflower (95.6%, 96.1%, 96.2% in cauliflower and 12%, 14%, 13.3% in cabbage accessions, respectively) (Fig. 4b). Same as *BoWUS2*, the highest expression of *BoMP2* and *BoMP3* was also observed at the curd formation stage (Fig. 4c).

#### Curd maintenance and floral arrest

meristems with floral meristems arrested in development. A large substitution (SV_b_70950) (~11.4 kb in OX-heart and ~7.7 kb in Korso) in the promoter region of the floral meristem identity (FMI) gene *BoCAL* was identified under strong selection. Almost all cauliflower (99.0%) and the majority of broccoli (87.5%) accessions shared the Korso allele, while most cabbage accessions (79.2%) harbored the OX-heart allele at this locus (Fig. 4b), suggesting its potential role in curd formation.

Several other FMI genes including *BoAP1.2, BoFUL1, BoFUL3* and *BoSEP3* were also affected by selected SVs (**Supplementary Table 9**), and all had relatively low expression at the vegetative, transition and curd formation stages, but significantly higher expression at the curd enlargement stage (**Supplementary Fig. 14**). Studies in *Arabidopsis* suggest that an antagonistic interaction between the inflorescence meristem identity (IMI) gene *TFL1* and FMI genes regulates the developmental fate transitions (Liljegren et al., 1999; Teo et al., 2014; Wagner, 2017). The nearly opposite expression pattern of *BoTFL1.2* compared to that of FMI genes indicated its repression role (**Supplementary Fig. 14**). While no selected SVs were identified in *BoTFL1.2*, a 13-bp deletion (SV_w_84836) in the intron of its positive regulator *BoAGL14* (Pérez-Ruiz et al., 2015) was found under strong selection (Fig. 4b), and *BoAGL14* showed the same expression pattern as *BoTFL1.2* (**Supplementary Fig. 14**), suggesting potentially important roles of both *BoTFL1.2* and *BoAGL14* in in floral identity arrest and inflorescence proliferation for curd formation and maintenance. SVP is a key negative regulator of floral transition (Hartmann et al., 2000; Gregis et al., 2013). A 420-bp (SV_w_74120) insertion in the promoter of *BoSVP1* was found in Korso and 98.1% of other cauliflower and all broccoli accessions, while only in 25.2% of the cabbage accessions (Fig. 4b). *BoSVP1* was significantly up-regulated from vegetative to transition stage, and kept high expression levels throughout the curd formation (Fig. 4c), indicating its repressor role in flower bud development, as reported in *Arabidopsis* (Liu et al., 2009). A cauliflower curd-specific gene, *BoCCE1*, was reported to have a potential role in the control of meristem development/arrest (Palmer et al., 2001; Duclos and Björkman, 2008). Here, we identified a 1,505-bp insertion (SV_b_67089a) covering the entire *BoCCE1* gene body in Korso. Genotyping of this insertion revealed that the *BoCCE1* gene was present in most cauliflower accessions (97.1%), but absent in most cabbage (86.5%) and broccoli (78.1%) accessions, suggesting a possible role of *BoCCE1* in floral arrest, as broccoli buds are arrested at later developmental stages compared to cauliflower buds (Fig. 4b,c).

#### Curd enlargement and spiral growth

Genes involved in organ size regulation, cell division and expansion, cell cycle etc. can regulate the curd weight. *CYP78A5 (KLU)* has been identified in *Arabidopsis* to prevent proliferation arrest and promote organ growth (Anastasiou et al., 2007; Stransfeld et al., 2010). The high expression of cauliflower *BoCYP78A5* was exclusively detected in curds (**Supplementary Table 9**), especially at the curd formation and enlargement stages (Fig. 4c). A 2775-bp substitution (SV_b_76292) in the promoter of *BoCYP78A5*, present in 98.1% of cauliflower accessions while only in 8.2% of cabbage accessions (Fig. 4b), might contribute to the curd-specific expression of *BoCYP78A5*. *BoARL2* (or *CDAG1*) has been proved to play a role in promotion of cauliflower curd size (Li et al., 2017). *BoARL2* was highly expressed in curd, with the highest expression during the curd enlargement phase of Korso (Fig. 4c). A 269-bp deletion (SV_w_38468) was detected in the promoter of *BoARL2*, and was present in all cauliflower and most broccoli (96.9%) accessions, while in only 41.7% of cabbage accessions (Fig. 4b).

The spiral arrangement of inflorescences is typical for cauliflower curds (Kieffer et al., 1998). Transcription of the *DRNL* gene marks lateral organ founder cells in the peripheral zone of the inflorescence meristem (Comelli et al., 2020). We identified a 258-bp deletion (SV_w_30645) in the promoter of *BoDRNL1*, which was present in all cauliflower and broccoli accessions while only in 40.6% of cabbage accessions (Fig. 4b). *BoDRNL1* was specifically expressed in the curd with the highest expression at curd formation and enlargement stages (Fig. 4c), implying its potential role in determining curd architecture. Selected SVs were also identified in the alphatubulin gene *BoTUA2* and four *BoTUA3* genes (Fig. 4b and **Supplementary Table 9**), whose homologue in *Arabidopsis* causes helical growth (Abe and Hashimoto, 2005).

## Discussion

The species *B. oleracea* includes a number of important vegetable crops displaying exceptionally high morphological diversity, with cauliflowers and cabbages representing two extreme morphotypes. In this study, we assembled high-quality chromosome-scale genome sequences for inbred lines of cauliflower and cabbage by integrating PacBio long-read sequences and Hi-C chromatin contact maps, which add important resources for future research and improvement of *B. oleracea* crops and provide the foundation for comprehensively exploring the phenotypic diversity of *B. oleracea*.

SVs play vital roles in the genetic regulation of plant phenotypic changes and are often the causative genetic variants for many important traits that are targets of crop domestication and breeding. However, population analysis of SVs in crops lags far behind that of SNPs, mainly due to the technological difficulties in accurate SV identification. The currently widely used SV calling approaches depend on the mapping of short sequencing reads to a reference genome, which are prone to both high false positive and high false negative rates (Mahmoud et al., 2019). The recent advances in long read sequencing technologies such as PacBio and Nanopore have helped read mapping in the detection of SVs. However, due to the restricted read length, some large SVs (e.g., insertions) cannot be detected (Alonge et al., 2020). In the present study, through direct comparison of the high-quality, reference-grade genome assemblies of cauliflower and cabbage, combined with long read mapping, we were able to identify ~120K high-confidence SVs, with a number of them larger than 100 kb. Genotyping of this reference set of SVs in a population comprising 271 accessions representing different *B. oleracea* morphotypes and investigation of allele frequency difference of these SVs in different morphotype populations, mainly cauliflower, cabbage and broccoli, revealed numerous SVs that are under selection in certain morphotypes, with many affecting genes associated with the corresponding unique phenotypes.

The curd of cauliflower is composed of thousands of inflorescence meristems that are spirally arranged on short enlarged inflorescence branches. This makes cauliflower an ideal model to analyze the genetic mechanism of inflorescence development and extreme organ genesis. SVs selected in cauliflower affected many genes. Combined with the analysis of expression profiles during curd development, we identified dozens of key SVs and associated genes that had potential associations with the unique curd phenotype of cauliflower. These included genes with roles in the different developmental stages of curd development. The first stage is curd initiation, involving the transition from the vegetative stage to the generative stage, with genes involved in floweringtime regulation affected (e.g., *FLC* and *FRI).* An essential step in curd formation is inflorescence proliferation, with genes like *WUS* and *MP* having cauliflower and broccoli-specific SVs. Cauliflower curds are further characterized by the floral meristem arrests, matching several floral identity genes (e.g., *CAL*, *AP1* and *SEP3)* as well as their potential negative regulatory genes (e.g., *AGL14, SVP* and *CCE1*) affected by selected SVs. Several genes with roles in cauliflower curd architecture were also affected by selected SVs. These include genes that play likely roles in organ size control (e.g., *CYP78A5* and *ARL)* and the curd spiral organization (e.g., *DRNL* and *TUA)* (Fig. 4d). Our analyses demonstrated the important contributions of SVs to the unique curd phenotype of cauliflower and shed light on the regulatory network of cauliflower curd formation.

## Methods

### Genome library construction and sequencing

Cauliflower (*B. oleracea* var. *botrytis*) accession Korso_1401 is a highly inbred line derived from Korso that was obtained from the Genebank of IPK Gatersleben (http://gbis.ipk-gatersleben.de; accession No. BRA2058), and has a white compact curd and long maturing time (> 95 d). Pointed cabbage (*B. oleracea* var. *capitate)* accession OX-heart_923, an inbred line with green pointed head and late bolting, was obtained from Vegetable Research Institute, Jiangsu Academy of Agricultural Science, Nanjing, China.

Young fresh leaves were collected from a single individual of each of the two accessions after a 24-h dark treatment and used for high molecular weight (HMW) DNA extraction using the cetyltrimethylammonium bromide method (Murray and Thompson, 1980). PacBio SMRTbell libraries were constructed from the HMW DNA using the SMRTbell Express Template Prep Kit 2.0 following the manufacturer’s protocols (PacBio). A total of 24 Single-Molecule Real-Time (SMRT) cells (9 from PacBio RSII and 15 from PacBio Sequel) for Korso and 15 SMRT cells (all from PacBio Sequel) for OX-heart were sequenced by NextOmics Biosciences Co., Ltd (Wuhan, China). For Illumina sequencing, paired-end libraries with insert sizes of ~400 bp were prepared using the NEBNext Ultra DNA Library Prep Kit and sequenced on a HiSeq 2500 system with 2×150 bp mode. Hi-C libraries were constructed using the Proximo Hi-C plant kit following the manufacturer’s instructions (Phase Genomics) and sequenced on an Illumina HiSeq X Ten system with 2×150 bp mode at Nextomics Biosciences.

For Korso, an optical map was generated using the Saphyr system (BioNano Genomics). Briefly, the HMW DNA labeling and staining were performed according to the manufacturer’s protocols, and then loaded onto chips and imaged on the Saphyr System according to the user guide. Data processing, construction of the Direct Label and Stain (DLS) optical maps and the hybrid map assembly were performed using the BioNano Genomics Access software suite.

For genome resequencing, young leaves from 163 different *B. oleracea* accessions (89 cauliflower, 65 cabbage and 9 broccoli accessions) were collected and used to extract DNA using the DNeasy Plant Mini Kit (Qiagen). Pair-end libraries with insert sizes of ~400 bp were constructed using the NEBNext Ultra DNA Library Prep Kit according to manufacturer’s instructions and sequenced on an Illumina HiSeq 2500 platform with the paired-end 2×150 bp (117 accessions) or 2×100 bp (46 accessions) mode.

### Transcriptome sequencing and data processing

Seven different tissues of Korso (root, stem, leaf, curd, bud, flower and silique) and OX-heart (root, stem, leaf, leafy head, bud, flower and silique) were collected for transcriptome sequencing. Roots, stems and leaves were sampled at the rosette stage of plants with 8-10 leaves (4-6 weeks after planting). The buds were about 2 mm in length, the flowers were blooming, and the siliques were at the developing stage including seeds. The curd of Korso and leafy head of OX-heart were collected when they were ready to be harvested. In addition, shoot apical meristem (SAM) samples from Korso were collected at the following developmental stages: vegetative, transition (curd initiation), curd formation (curd diameter of ~1 cm), pre-mature (curd diameter of 10 cm), and branch elongation (mature). Two or three independent biological replicates were performed for each sample. RNA was extracted from each tissue using the TIANGEN RNAprep Pure Kit (Cat: No. DP441). RNA quality was assessed using an Agilent 2100 BioAnalyzer.

RNA-Seq libraries were prepared using the Illumina TruSeq RNA Sample Prep Kit and sequenced on an Illumina HiSeq X Ten system at WuXi NextCODE (Shanghai, China). Raw RNA-Seq data were preprocessed using the NGS QC Toolkit (v2.3.3) (Patel and Jain, 2012) to remove adaptors, low quality bases, and reads containing more than 10% unknown bases (‘N’). The cleaned reads were mapped to the reference genomes allowing up to two mismatches using HISAT2 (Kim et al., 2019). Based on the alignments gene expression levels were estimated as fragments per kilobase of transcript per million mapped fragments (FPKM). Differential expression analysis between different developmental stages of SAM samples was performed using the DESeq package (Anders and Huber, 2010).

The PacBio Iso-Seq library was also constructed for Korso using equally mixed RNA samples from the seven tissues. The cDNA synthesis and amplification were performed using the NEBnext Single Cell/Low Input cDNA Sythesis & Amplification Module kit. SMRTbell libraries were constructed with the SMRTbell Express Template Prep kit 2.0. Three SMRT cells were sequenced on the PacBio Sequel system with the Sequel DNA Polymerase 2.0 and Sequel Sequencing Plate 2.0 by NextOmics Biosciences Co., Ltd (Wuhan, China). Different subreads from the same polymerase read were used to generate circular consensus sequences (CCSs). The CCSs were then classified as full-length, non-chimeric and non-full-length according to the presence or absence of 5′-primer, 3′-primer, and poly A/T tails. These sequences were then clustered using an Iterative Clustering and Error (ICE) correction algorithm incorporated in the IsoSeq_SA3nUP pipeline (https://github.com/PacificBiosciences/IsoSeq_SA3nUP). The resulting consensus isoforms were first polished using the non-full-length reads and then the RNA-Seq reads with LoRDEC (v0.3) (Salmela and Rivals, 2014).

### Genome size estimation

The genome sizes of Korso and OX-heart were estimated using flow cytometry. For each sample, ~1g leaves were finely chopped with a razor blade in 2000 μl LB01 isolation buffer (Doležel et al., 1989). The resulting suspension was filtered through 30-μm nylon, and then 2 μl 10 mg ml^−1^ RNase I was added in room temperature for 15 min. After centrifugation at 1000r/min for 5min, the supernatant was discarded, and the precipitated nuclei were collected. The nuclear DNA was fluorescently labeled with PI (Propidium iodide) staining solution and stained in the dark for 30 minutes. The DNA peak ratio was assessed by flow cytometry (BD FACSCalibur system, BD Biosciences) using *B. rapa* as the internal reference. The ModFit software (Verity Software House) was used for data analyses.

### *De novo* genome assembly

PacBio reads were *de novo* assembled using Falcon (v1.8.7) (Chin et al., 2016). The assembled contigs were first polished with Arrow (Chin et al., 2013) using PacBio long reads, and then further polished with Illumina shotgun reads using Pilon (Walker et al., 2014). The polished Korso contigs was further improved by BioNano optical genome maps. The PacBio contigs were anchored to optical maps to construct scaffolds and the resulting gaps between connected contigs were filled using PBJelly (https://github.com/esrice/PBJelly) with the following parameters: ‘-minMatch 8 -sdpTupleSize 8 -minPctIdentity 75 -bestn 1 -nCandidates 10 -maxScore -500 -noSplitSubreads’. The resulted contigs were polished again by Arrow and Pilon as described above.

To remove potential contaminations in the assemblies, the final contigs were divided into 50-kb bins and then searched against the GenBank nucleotide (nt) database using blastn (Camacho et al., 2009). Sequences with best hits not in the green plants were possible contaminations and discarded.

To construct pseudomolecules, raw Hi-C reads were processed to trim adapters and low-quality sequences using Trimmomatic (Bolger et al., 2014) with parameters ‘SLIDINGWINDOW:4:20 MINLEN:50’. The cleaned reads were aligned to the final contigs using bowtie2 end-to-end algorithm (Langmead and Salzberg, 2012). HiC-Pro pipeline (Servant et al., 2015) was then used to remove duplicated read pairs, detect valid ligation products and perform quality controls. The assembled contigs were then clustered, ordered, and oriented into pseudomolecules using Lachesis (Burton et al., 2013).

### Repetitive element and centromere prediction

A custom repeat library was constructed for each genome according to the pipeline described in http://weatherby.genetics.utah.edu/MAKER/wiki/index.php/Repeat_Library_Construction-Advanced, using MITE-Hunter (Han and Wessler, 2010), RepeatModeler (http://www.repeatmasker.org/RepeatModeler/), and LTRharvest and LTRdigest from GenomeTools (Gremme et al., 2013). Repeat sequences were identified by scanning each genome assembly using the corresponding repeat library with RepeatMasker (http://www.repeatmasker.org/).

Full-length long terminal repeat retrotransposons (LTR-RTs) were identified using LTR_finder (Xu and Wang, 2007) and LTR_harvest (Ellinghaus et al., 2008), and redundancies in the full-length LTR-RTs were then removed using LTR_retriver (Ou and Jiang, 2018). The substitution rate between the two end sequences of each LTR-RT was calculated using PAML (Yang, 2007). The LTR expansion time was estimated according to the formula T=S/2μ, where S is the substitution rate and μ is the mutation rate (1.5×10^-8^ per site per year) (Koch et al., 2000).

The previously reported centromeric satellite repeats, including CentBr, CRB, TR238 and PCRBr (Lim et al., 2007), were used to scan the nine chromosomes of both Korso and OX-heart genome assemblies. The locations of centromeres were estimated based on the peak regions of these centromeric satellite repeats.

### Prediction and annotation of protein-coding genes

Protein-coding genes were predicted from Korso and OX-heart assemblies using EVidenceModeler (EVM) (Haas et al., 2008) by integrating transcript evidence, *ab initio* prediction and protein homology searching. For transcript evidence, the Illumina RNA-Seq reads were mapped to the reference genomes using HISAT2 (Kim et al., 2019) and assembled into transcripts using StringTie (Pertea et al., 2015). PASA (Haas et al., 2008) was then used to determine potential intron-exon boundaries. The PacBio Iso-Seq reads of Korso were also used as the transcript evidence. For *ab initio* prediction, AUGUSTUS (Stanke and Morgenstern, 2005), SNAP (Korf, 2004), and GENSCAN (Burge and Karlin, 1997) were employed. AUGUSTUS and SNAP were trained for each genome using the high-confident gene models obtained with the PASA analysis, while the *Arabidopsis* gene models were used to train GENSCAN. For protein homology searching, protein sequences from relative and model species (*Arabidopsis thaliana*, *Aethionema arabicum*, *Brassica napus*, *Brassica nigra*, *Brassica rapa*, *Capsella rubella*, *Thellungiella halophilla*, *Brassica oleracea* TO1000 and HDEM) were aligned to the genome assemblies using GenBlastA (She et al., 2009) and based on the alignments GeneWise (Birney et al., 2004) was then used to predict gene structures. Finally, EVM was used to generate a consensus gene set for each genome by integrating evidence from transcript mapping, protein homology and *ab initio* predictions. To annotate the predicted protein-coding genes, their protein sequences were searched against GenBank nr, SwissProt, KEGG and TrEMBL protein databases. GO term annotation and enrichment analysis were carried out using the Blast2GO suite (Gotz et al., 2008).

### SV identification between Korso and OX-heart genomes

To identity SVs between genomes of OX-heart cabbage and Korso cauliflower, we first aligned the two genomes using the Minimap2 with parameters ‘-ax asm5′ (Li, 2018). The resulting alignments were analyzed using Assemblytics (Nattestad and Schatz, 2016) to call SVs. The resulting SVs spanning or close (distance <50bp) to gap regions in either of the two genomes were removed.

We further identified SVs by aligning PacBio reads from OX-heart to the Korso genome and Korso PacBio reads to the OX-heart genome, using Minimap2 with parameters: ‘-eqx -L -O 5,56 -e 4,1 -B 5 --secondary=no -z 400,50 -r 2k -Y --MD -ax map-pb’. Based on the alignments, SVs were called using pbsv (https://github.com/PacificBiosciences/pbsv). The identified SVs spanning gap regions in the genomes were discarded. To further evaluate the SVs, sequences of the two 5-kb flanking regions of each SV were extracted from the genome and then blasted against to the other genome. The unique alignments between the two genomes identified by Assemblytics were used to filter SVs identified by pbsv. SVs were kept if the reliable blast hits of the two flank sequences (alignment length >50 bp, identity >90% and e-value <1e-10) were found in the expected region on the query genome, and the gap size between the two hits was consistent with that estimated by pbsv. Specifically, for insertions, we required that the deviation between pbsv estimated size and the distance observed between the two blast hits was smaller than 20%. For deletions, the allowed gap or overlap between the two blast hits of flanking regions should be smaller than 3 bp.

It is noteworthy that, besides the simple indels with defined breakpoints, Assemblytics also reported four types of complex SVs without defined breakpoints, including repeat expansion, repeat contraction, tandem expansion and tandem contraction. If SVs identified by pbsv were in the regions of these complex SVs, their precise breakpoints could be defined, i.e., these complex SVs could be converted into one or more simple indels. SVs identified by Assemblytics and pbsv were merged if they overlapped with each other by at least 50% of their lengths.

### Genotyping of SVs in *B. oleracea* accessions

Raw genome sequencing reads of the 163 *B. oleracea* accessions generated in this study and 108 accessions reported previously (Cheng et al., 2016) were first processed to consolidated duplicated read pairs into unique read pairs. Duplicated read pairs were defined as those having identical bases in the first 90 bp for 100-bp reads or 100 bp for 150-bp reads of both left and right reads. The resulting reads were then processed to trim adapters and low-quality sequences using Trimmomatic (Bolger et al., 2014) with parameters ‘SLIDINGWINDOW:4:20 MINLEN:50’.

To genotype SVs in these accessions, the cleaned reads were aligned to the OX-heart and Korso genomes, respectively, using BWA-MEM (Li, 2013), allowing no more than 3% mismatches. For each SV in each accession, we checked reads aligned to the regions spanning the breakpoints of the SV in both OX-heart and Korso genomes. For each breakpoint, we first required at least 3 split reads to support the SV call. If there were no enough split reads supporting the SV, we then checked the read coverage in the SV region. For a deletion, we required that <50% of the deleted region was covered by reads with 2× depth, while >50% of at least one flanking region with the same length of the deleted region was covered. Based on the split read and read depth information, SVs in a particular accession could be classified as Ox-heart genotype (same genotype as Ox-heart), Korso genotype (same genotype as Korso), heterozygous (containing both Ox-heart and Korso genotypes) and undetermined (genotype that were not be able to determined due to insufficient read mapping information).

For population analyses, we divided the genome into 25-kb non-overlapping windows and randomly selected one SV per window. The genotype data of the chosen SVs of the entire *B. oleracea* population were used to construct a maximum-likelihood phylogenetic tree using IQ-TREE (Nguyen et al., 2015) with 1,000 bootstraps. The same SVs were also used to perform the principal component analysis (PCA) with TASSEL5 (Bradbury et al., 2007) and to investigate population structure of *B. oleracea* accessions with STRUCTURE (Hubisz et al., 2009). Allele frequencies of 84,571 SVs with its genotype determined in at least 50% of accessions in both cauliflower and cabbage populations were calculated. Significance of the difference of the SV allele frequencies between cauliflower and cabbage groups was determined using Fisher’s exact test, and the resulting raw *P* values were corrected using the Bonferroni method. SVs with adjusted *P* values <0.001 and fold change ≥ 2 were defined as those highly differentiated between cauliflower and cabbage.

### Data Availability

The genome assemblies and raw genome and transcriptome sequencing reads of Korso and OXheart have been deposited into the NCBI BioProject database (http://www.ncbi.nlm.nih.gov/bioproject) under accession numbers PRJNA546441 and PRJNA548819, respectively.

## ACKNOWLEDGMENTS

We thank Dr. Evelyn Klocke for providing the original seeds of Korso. This research was supported by grants from the National Key R&D Program of China (2016YFD0100204-14 to F.L. and 2016YFD0101702 to Jianbin Li), President Foundation for Science of Beijing Academy of Agricultural and Forestry Sciences (YZJJ201701 to F.L.), International Cooperation Foundation of Beijing Academy of Agricultural and Forestry Sciences (GJHZ2017-2 to F.L. and N.G.), National Modern Agriculture Industry Technology System (CARS-25 to Jianbin Li), Innovation Program of Beijing Academy of Agricultural and Forestry Sciences (KJCX20200205 to F.L. and KJCX20200108 to N.G.), National Natural Science Foundation of China (32072576 to N.G.). Overseas Training Program of State Administration of Foreign Experts Affairs of China (P172001038 to N.G.), and the U.S. National Science Foundation (IOS-1855585 to Z.F.).

## AUTHOR CONTRIBUTIONS

F.L., Z.F., Jianbin Li, G.B. and N.G. designed and managed the project. Jianbin Li., S.W., C.Z., H.J., G.B., S.H., M.Z., F.Y. and W.Z. collected samples. Jingjing Li, M.Y. and Y.L. contributed to the Korso and OX-heart genome assemblies and annotations. M.D., G.W., L.M. and N.G. generated RNA-Seq data. L.G., Y.L., H.S. and N.G. performed SV calling and genotyping. L.G., N.G., Y.L., Y.P., T.B. and H.S. performed data analysis. N.G. and L.G. wrote the manuscript. Z.F., F.L., G.B. and Y.X. revised the manuscript.

## COMPETING INTERESTS

The authors declare no competing interests.

## Notes

### Competing Interest Statement

The authors have declared no competing interest.

